# Neural dynamics of confidence formation under uncertainty

**DOI:** 10.64898/2026.09.08.750275

**Authors:** Tomoya Okaguchi, Kazumasa Uehara

## Abstract

Our perceptual decision-making can vary depending not only on changes in the surrounding environment but also on metacognitive processes. In particular, the balance between confidence and uncertainty may play a critical role in shaping behavior and the underlying neural dynamics over time, yet how this balance operates remains largely unclear, especially when uncertainty is high and confidence is low, or vice versa, during perceptual decision-making.

To address this, we employed the Multi-Attribute Attention Task, along with an option to opt-out of perceptual judgments. During this task, we recorded multichannel electroencephalography (EEG) data from 26 healthy individuals. Our results showed that EEG-based functional connectivity increased in density across the whole brain, which was associated with the emergence of confidence during perceptual decision-making. Moreover, increased functional connectivity density emerged in correct trials but was not observed in incorrect trials. Notably, this density increase was specific to correct trials and became evident prior to the decision being made. Thus, correct decisions may be characterized by the early recruitment of distributed neural networks that support the integration of sensory evidence and the formation of reliable confidence.

Building on these findings, we provide novel evidence that confidence formation under uncertainty relies on dynamic changes in densely interconnected networks across distributed brain regions that support highly accurate and high-confidence perceptual decision-making in humans.

## Introduction

In daily life, we are frequently required to make decisions based on incomplete or ambiguous sensory information as well as an uncertain and volatile surrounding environment, such as judging the safety of crossing a street in dense fog, to high-stakes scenarios in team sports, where players must predict the movement of a ball or opponent players using vision and auditory stimuli. Decisions made in environments where information about the current state or potential outcome is insufficient are categorized as decision-making under uncertainty. In these instances, rather than relying solely on insufficient sensory input, our central nervous system frequently depends on an internal, subjective evaluation known as confidence (Grimaldi et al., 2015). In fact, confidence has been conceptualized as a core component of metacognition, which refers to the capacity to monitor and evaluate one’s own cognitive processes (Boldt et al., 2019; Fleming & Lau, 2014). Given that these cognitive processes are subjective, identifying a distinct neural process within the brain remains a pivotal challenge in the field of cognitive neuroscience.

To address this open question, current studies on perceptual decision-making have identified several neurophysiological correlates of confidence. Electroencephalography (EEG) studies in humans have highlighted event-related potentials (ERPs) arising from the centro-parietal regions, most notably the Centro-Parietal Positivity (CPP), which reflects the accumulation of sensory evidence (Herding et al., 2019), and the Contingent Negative Variation (CNV), which represents neural preparation modulated by task difficulty (Boldt et al., 2019). Likewise, functional magnetic resonance imaging (fMRI) studies have anatomically localized confidence-related processing to a distributed brain network, including the striatum, dorsomedial prefrontal cortex (dmPFC), cingulate cortex, insula, and other regions within the frontal, parietal, and occipital cortices (Grimaldi et al., 2015; Paul et al., 2015).

Nevertheless, some critical gaps remain in the understanding of confidence-related neural representations. First, a substantial temporal gap seems to exist between the decision-making process and the measurement of confidence. In principle, traditional paradigms for investigating confidence during behavior rely on post-decisional self-reports, which may suffer from decay or distortion of metacognitive information between action and the subsequent self-report of confidence, typically expressed on a numerical scale (Fleming et al., 2018). As alternatives, behavioral methods that infer confidence from action-related states without explicit introspective reporting, such as Post-Decision Wagering (PDW) (Persaud et al., 2007) and the “Opt-out” paradigm (Kiani and Shadlen, 2009), have been proposed as useful protocols. These approaches allow declining a trial to avoid potential errors, thereby enabling a more direct and natural estimation of latent confidence. However, the temporal dynamics of neural activity related to latent confidence remain not fully understood.

In addition, the functional relationships between the identified brain regions, including their temporal dynamics, have not been fully investigated. Neural dynamics on the millisecond timescale are thought to be essential for decision-making (Demetriou et al., 2018), and confidence, much like the decision process itself, is believed to rely on the dynamic accumulation of sensory-cognitive information (Gherman & Philiastides, 2018). To bridge this gap, neural recordings with superior temporal resolution are required to characterize changes in functional connectivity between local and distributed cortical areas as a function of decision-making.

Moreover, real-world decision-making rarely occurs under fully specified conditions. Consequently, individuals are often required to act under contextual uncertainty, in which relevant information is incomplete, ambiguous, or dynamically changes with the surrounding environment. Incorporating contextual uncertainty into decision-making experiments is therefore essential for capturing the cognitive and neural processes that support adaptive changes in metacognition in naturalistic environments. We assume that contextual uncertainty modulates the accumulation of sensory evidence, the weighting of prior knowledge, and the formation of confidence, thereby governing both behavioral choices and subsequent execution. By explicitly manipulating contextual uncertainty, experimental paradigms can dissociate decision-related processes from primary sensory processing and reveal how the brain flexibly reorganizes its neural architecture in response to changing environmental demands. Such approaches offer critical insights into the neural mechanisms underlying the interplay between decision-making and confidence under uncertainty.

Here, the goal of the present study was to examine how contextual uncertainty affects confidence and perceptual decision-making at the behavioral level and to characterize the underlying neural dynamics. To do so, we leveraged multichannel scalp EEG recordings to examine the temporal neural dynamics of confidence during perceptual decision-making under varying levels of uncertainty. Furthermore, an experimental task incorporating an opt-out paradigm enabled the separation of confidence-related neural activity from primary perceptual processing. This experimental paradigm allows us to unmask neural mechanisms underlying not only uncertainty-related decision-making but also quitting subsequent action, i.e., opting out, caused by high uncertainty. We hypothesize that contextual uncertainty would modulate confidence and functional brain connections for perceptual decision-making. In particular, functional connectivity density will differentiate high-confidence from low-confidence decisions. Increased functional connectivity density may reflect enhanced communication and integration among distributed neural systems supporting confident perceptual decisions.

## Methods

### Participants

All participants were naïve to the purpose of the study. Twenty-six healthy adults (mean age = 22.19 years, standard deviation (SD) = 1.21; 2 females, 3 left-handed) participated in the experiment. Their handedness was assessed using the Edinburgh Handedness Inventory (Oldfield, 1971). Participants with any history of neurological or psychiatric disorders were excluded. The study was approved by the Institutional Review Board for Human Research at Toyohashi University of Technology (Approval No. 2023-06) in accordance with the guidelines established in the Declaration of Helsinki. All participants provided written informed consent before the data collection.

### Experimental task and protocols

The data collection was conducted in a dark, sound-attenuated room. Participants sat upright on a comfortable chair. To display visual stimuli throughout the data collection, we used a 24.5-inch PC monitor (XL2546K-B, BenQ; refresh rate of 144 Hz), placed in front of a participant. This monitor was positioned at 50 cm from the chin rest, aligned with the participant’s line of sight. Their heads were stabilized using a chinrest (SR-HDR, SR Research Ltd.). The experiment began with a 6-minute resting-state EEG recording (3 min eyes open, 3 min eyes closed). During this recording, the eyes-closed condition required maintaining a resting state for 3 min without any thoughts or body movement. In the eyes-open condition, subjects were instructed to view video footage of natural scenery for 3 min without any thoughts or body movement. Before beginning the main task, participants completed the familiarization session consisting of 40 trials. Subsequently, participants were required to complete the main task.

### Experimental Task

The behavioral task used in the present study was inspired by the study of Kosciessa et al. (2021) (Kosciessa et al., 2021). Drawing on uncertainty-manipulation approaches, referred to as the Multi-Attribute Attention Task, we examined both uncertainty-related decision-making and opt-out choices. In other words, we sought to characterize how the temporal dynamics of neural activity change as a function of the balance between confidence and uncertainty. The task was developed and implemented in Python (ver. 3.13.0) using the PsychoPy library (Peirce et al., 2019). A trial structure is illustrated in Figure 1A. Each trial began with a 3-second cue period during which a minimum of one and a maximum of four visual attributes were presented on the PC monitor.

**Figure 1:**
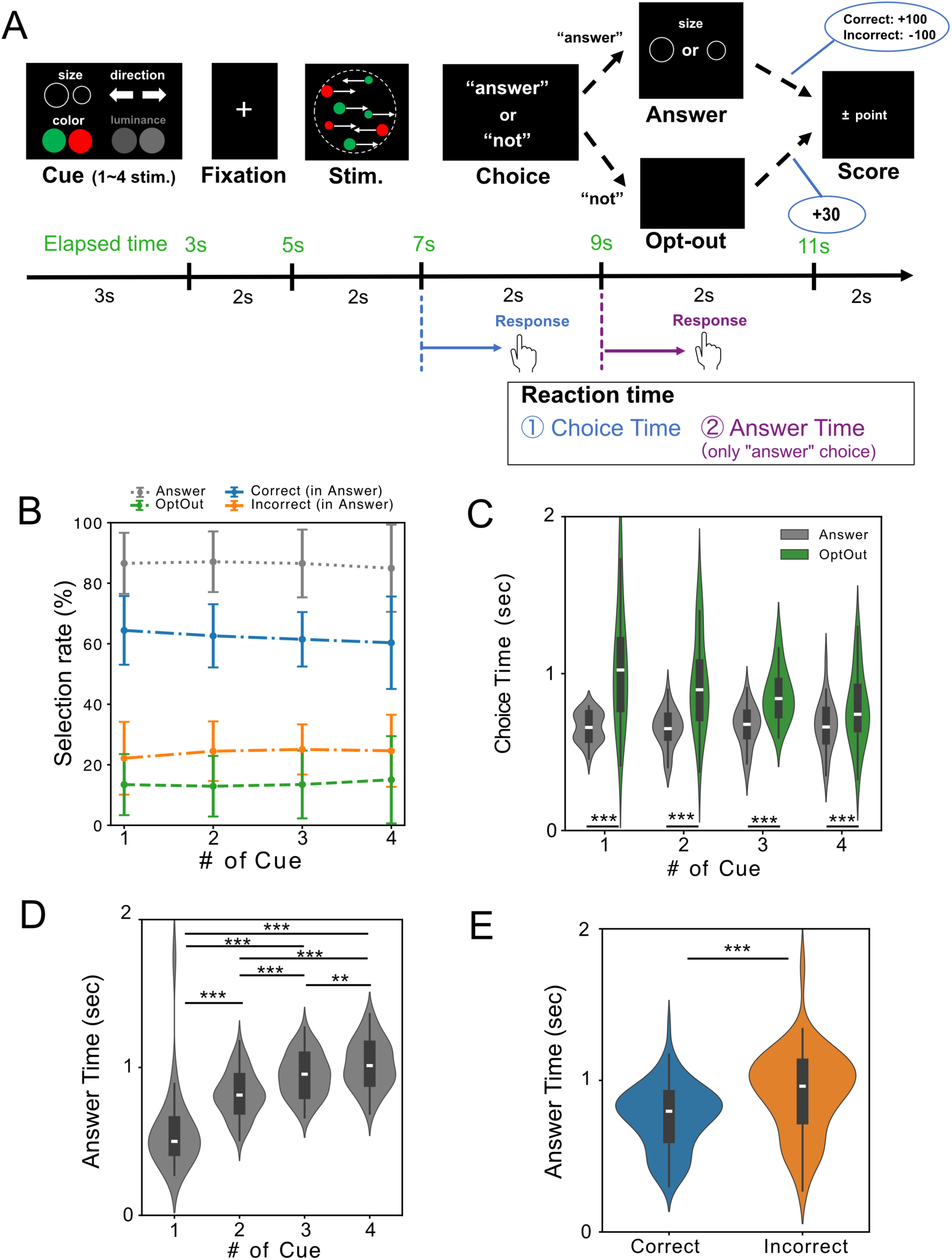
Trial structure and behavioral findings. **(A)** Each trial began with a 3-second cue period during which a minimum of one and a maximum of four visual attributes were presented on the PC monitor. The provided visual attributes consisted of random moving dots that varied simultaneously across four attributes: motion direction (left/right), color (green/red), size (large/small), and brightness (light/dark). Using these randomized visual attributes, we manipulated uncertainty in decision-making. Next, after a 2-second fixation period, which displayed a fixation cross on the center of the monitor, a multi-attribute random-dot stimulus was presented for 2 seconds. Two seconds were then given to make a metacognitive choice: to “answer” (challenge the task) or “not answer” (opt-out). During this period, participants were required to press either the left or right arrow key on a PC keyboard, corresponding to “answer” and “not answer” options, respectively. If they chose to “answer”, another 2-second time window was given to report the dominant feature of the one, along with cued attributes. To maintain participants’ motivation for confidence-based choices, a correct answer was rewarded with +100 points on each trial, an incorrect answer was penalized with -100 points, and selecting the avoidance option yielded a fixed gain of +30 points. **(B)** Percentage of behavioral choices across all trials as a function of the number of cues. Multichannel scalp EEG data were continuously recorded throughout the experiment. **(C)** Choice reaction time (Choice time) for “Answer” and “Opt-out” responses across all trials as a function of the number of cues. **(D)** Answer reaction time (Answer time) for responding to a question concerning the dot’s feature as a function of the number of cues. **(E)** Comparison of Answer reaction time (Answer time) between the correct and incorrect trials.

The provided visual attributes consisted of random moving dots that varied simultaneously across four attributes: motion direction (left/right), color (green/red), size (large/small), and brightness (light/dark). Both the number of attributes and their combinations were randomized across trials. Using these randomized visual attributes, we manipulated uncertainty in decision-making. We hypothesized that a reduction in uncertainty would result in higher decision confidence among participants. Next, after a 2-second fixation period, which displayed a fixation cross on the center of the monitor, a multi-attribute random-dot stimulus was presented for 2 seconds. Two seconds were then given to make a metacognitive choice: to “answer” (challenge the perceptual task) or “not answer” (quit the task). During this period, participants were required to press either the left or right arrow key on a PC keyboard, corresponding to “Answer” and “Not (i.e., Opt-out)” options, respectively. If they chose to “Answer”, another 2-second time window was given to report the dominant feature of the one, along with cued attributes. To maintain participants’ motivation for confidence-based choices, a correct answer was rewarded with +100 points on each trial, an incorrect answer was penalized with -100 points, and selecting the avoidance option yielded a fixed gain of +30 points. Every eight trials, participants received visual feedback on their overall performance via the cumulative score displayed on the PC monitor. Note that task parameters as well as calibration of task difficulty involving cognitive efforts were determined based on a preliminary pilot study (N=4; they did not participate in the main experiment). Based on this pilot testing, we confirmed that our uncertainty manipulation was effective and finalized the incentive structure (e.g., +30 points for avoidance). Additionally, the difficulty for each attribute was individually calibrated by adjusting the proportion of dots favoring the dominant feature (e.g., 55% right-moving vs. 45% left-moving dots). This calibration was based on each participant’s training performance and aimed to achieve an average accuracy of approximately 60% in the main experiment. The main experiment consisted of 256 trials, divided into four blocks of 64 trials, with short breaks between the blocks.

### Data acquisition

#### Behavioral data

On each trial, reaction times (RTs) for both the decision to either answer or opt-out of the trial and the subsequent perceptual response to the random moving dots were collected using the PC keyboard.

#### EEG recording

To record neural oscillations throughout the experimental task, we used a multi-channel EEG amplifier system (actiCHamp Plus, Brain Products, Ltd). EEG data were continuously recorded from 63 scalp electrodes positioned according to the layout of the international 10-10 system using active electrodes embedded in a wearable elastic cap (actiCAP). The ground electrode was placed at AFz, and the left mastoid served as the recording reference. Four EOG channels were used to monitor horizontal and vertical eye movements. The EEG and EOG signals were amplified, digitized with 24-bit resolution, and sampled at 1000 Hz. Event triggers generated via PsychoPy were integrated into the EEG and EOG recordings in real time. Throughout the data collection, skin/electrode impedance was kept below 10kΩ.

### Data Analysis

#### EEG data

EEG data were preprocessed using the MNE-Python library (Gramfort, 2013). With regard to preprocessing and artifact removal, the following preprocessing steps were performed: (1) Re-referencing the data to the average of the left and right mastoids. (2) Applying a 1-60 Hz bandpass filter and a 60 Hz notch filter. (3) Removing artifacts caused by eye movements, blinking, and muscle contraction using Independent Component Analysis (ICA). (4) Epoching the data relative to event onsets. (5) Rejecting epochs with amplitudes exceeding ±100 µV. (6) Applying a Current Source Density (CSD) transformation to reduce volume conduction effects and improve spatial precision (F. Perrin et al., 1987). The preprocessed EEG data were downsampled from 1000 to 250 Hz and then analyzed using the following post-processing procedures.

To quantify cognitive processes, we utilized a well-known traditional electrophysiological measure, namely event-related potentials (ERPs). In the present study, we computed the ERPs by averaging across trials within each condition and aligned them to the onset of the random dot motion, which occurred 5 s after trial onset (see Figure 1A). We particularly focused on changes in ERPs-labeled centroparietal positivity (CPP) appears to trace evidence accumulation during decision making (Herding et al., 2019; Ko et al., 2024; O’Connell et al., 2012). This metric provides evidence for the validity of our experimental task by demonstrating that manipulating the number of cues successfully modulated perceptual uncertainty. Moreover, to investigate how decision-making under uncertainty modulates neural oscillations and cortical neural coupling, time-frequency decomposition was performed using Morlet wavelets (Van De Vijver & Cohen, 2019), with frequencies ranging from 1 to 45 Hz in 1 Hz-steps. To optimize the trade-off between time resolution and frequency resolution, the number of wavelet cycles varied from 3 to 14, increasing with frequency. The complex wavelet transform is expressed by the following equation:

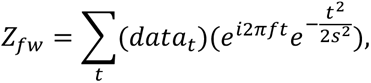

where *s* = *n*/2*πf*. Here, *f* is the frequency, *w* is the time window, *f* represents the time points in *w*, and *data_t_* is the signal at time *t*. Additionally, functional connectivity was assessed by computing the weighted Phase Lag Index (wPLI) (Vinck et al., 2011) from the complex-valued time-frequency data. The wPLI extends the phase lag index (PLI) and offers several advantages over it. Specifically, the wPLI is robust to sample-size bias and noise and provides greater statistical power to detect changes in phase synchronization. Given these advantages, the wPLI was employed as an index to characterize functional brain networks in the present study. The wPLI is defined as:

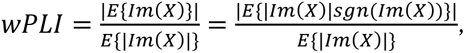

where *E*{⋅} is the expected value, *sgn*(⋅)is the sign function, *Im*(⋅)is the imaginary part of a complex number, and *X* is the cross-spectral density between two signals. To account for sample-size bias, we employed the debiased WPLI-square estimator (Vinck et al., 2011). This process yielded a 62x62 adjacency matrix for each trial, which was then thresholded to create binarized graphs for further analysis. The threshold for determining functionally meaningful network pathways was set to the 0.5th percentile value of wPLI within each period.

### Statistics

#### Behavioral data

To examine the effect of cue number (number of cues: 4 levels) on choice behavior, we fit binomial GLMMs (logit link; lme4::glmer, R software, version 4.4.1), decomposing the three-way outcome into two binary models: Model 1 (Opt-Out vs. Answer) and Model 2 (Correct vs. Incorrect, Answer trials only). The number of cues was treated as an ordered fixed factor, with Subject as a random effect (maximal random-slope structure retained if it converged; otherwise, random intercepts only). The overall effects of the number of cues were assessed via a likelihood ratio test against a reduced model. Reaction times were separately analyzed with two-way repeated-measures ANOVAs using ANOVAKUN (ver. 4.8.9) implemented on R software. In addition, Answer RT (Answer trials only) was analyzed with number of cues × response accuracy (Correct vs. Incorrect), and Challenge RT was analyzed with number of cues × response type (Answer vs. Opt-Out). Degrees of freedom were adjusted by the Chi-Muller epsilon correction for sphericity violations. If a significant main effect was detected (p < 0.05), post hoc pairwise comparisons were conducted using paired t-tests with Shaffer’s modified sequentially rejective Bonferroni procedure.

#### EEG data

For ERP data, we used a Wilcoxon signed-rank test at each time-electrode pair, with False Discovery Rate (FDR) correction for multiple comparisons. A cluster-based permutation paired t-test was applied to assess differences between the opt-out and answer conditions and between the correct and incorrect conditions, using two-tailed testing with 1,000 random permutations and family-wise error rate (FWER) correction applied at the cluster level. Statistical significance was corrected for multiple comparisons across electrodes, time points, and frequencies for each test.

For EEG channel-based functional connectivity estimated by wPLI, differences in wPLI across conditions within each network were assessed using the Network-Based Statistic (Zalesky et al., 2010). For each edge, paired t-tests were first conducted between the conditions. Suprathreshold edges (p < 0.001) were then determined and assembled into connected components. Statistical significance of component size was determined using 5,000 random permutations, with FWER correction applied at the network level. Unless noted otherwise, components with FWER-corrected p < 0.05 were considered statistically significant.

## Results

### Behavioral responses

Figure 1B shows the percentage of behavioral choices across all trials according to the number of cues. Although there were no significant differences in the probability of behavioral choices across the number of cues (GLMM (Accuracy): χ² (3) = 2.6409, p = 0.4504), reaction times for Answer (i.e., Answer time) increased significantly with the number of cues (Figure 1D) (ANOVA: F_3, 75_ = 205.81, p = <0.001). Although our manipulation approach, namely the number of visual cues, did not strongly affect participants’ behavioral choices, the results of answer time indicate that the manipulation task used in the present study can induce perceptual uncertainty.

Figure 1C displays reaction times according to whether participants selected the answer or opt-out option after viewing the random moving dots. Repeated-measures ANOVA with the factors number of cues and choice (challenge vs. opt-out) revealed significant main effects of number of cues: F_3, 75_ = 3.05, p = 0.033) and choice (F_1, 25_ = 250.9, p < 0.001). A significant interaction between the two factors was also observed (F_3, 75_ = 3.70, p = 0.015). Follow-up analyses showed that the main effect of number of cues was significant only in the opt-out condition (number of cues in the Answer condition: F_3, 75_ = 0.80, p = 0.492, number of cues in the opt-out condition: F_3, 75_ = 2.94, p = 0.038). However, post hoc pairwise comparisons did not reveal any significant differences between the numbers of cues within the opt-out condition. In addition, reaction times were significantly longer in the opt-out trial than in the Answer trial at every cue level (all p < 0.05). These findings suggest that opt-out required greater deliberation than accepting the challenge, and that this deliberation was selectively influenced by the amount of available cue information. Figure 1D shows the Answer time as a function of the number of cues and response accuracy (correct vs. incorrect). Repeated-measures ANOVA with the factors number of cues and correct/incorrect revealed significant main effects (number of cues: F_3, 75_ = 205.80, p < 0.001, correct/incorrect: F_1, 25_ = 155.59, p < 0.001) but no significant interaction between the two factors (F_3, 75_ = 1.85, p = 0.145). Post hoc pairwise comparisons showed that Answer time was significantly increased as the number of cues increased (all p < 0.01). Moreover, Answer times were significantly shorter for correct responses than for incorrect responses (p < 0.001). Taken together, perceptual uncertainty affected the time required to reach a decision, rather than the decision itself, and longer reaction times were associated with both opting out and incorrect responses. These findings imply that reaction time may capture internal decision dynamics that are not reflected in overt behavioral choices.

#### EEG data

Regarding changes in ERP components during decision-making, cluster-based permutation testing identified significant two CPP components (p = 0.0006). These components were significantly greater in the Answer trials than in the Opt-out trials approximately 5.5 s after cue onset. (i.e., 0.5 s after the onset of the random dot motion) (p = 0.002, after FWER correction) (Figure 2). This indicates that a larger amount of decision evidence had been accumulated before participants committed to a perceptual judgment. In contrast, reduced CPPs in opt-out trials indicate that insufficient evidence accumulation preceded the decision to opt-out.

**Figure 2:**
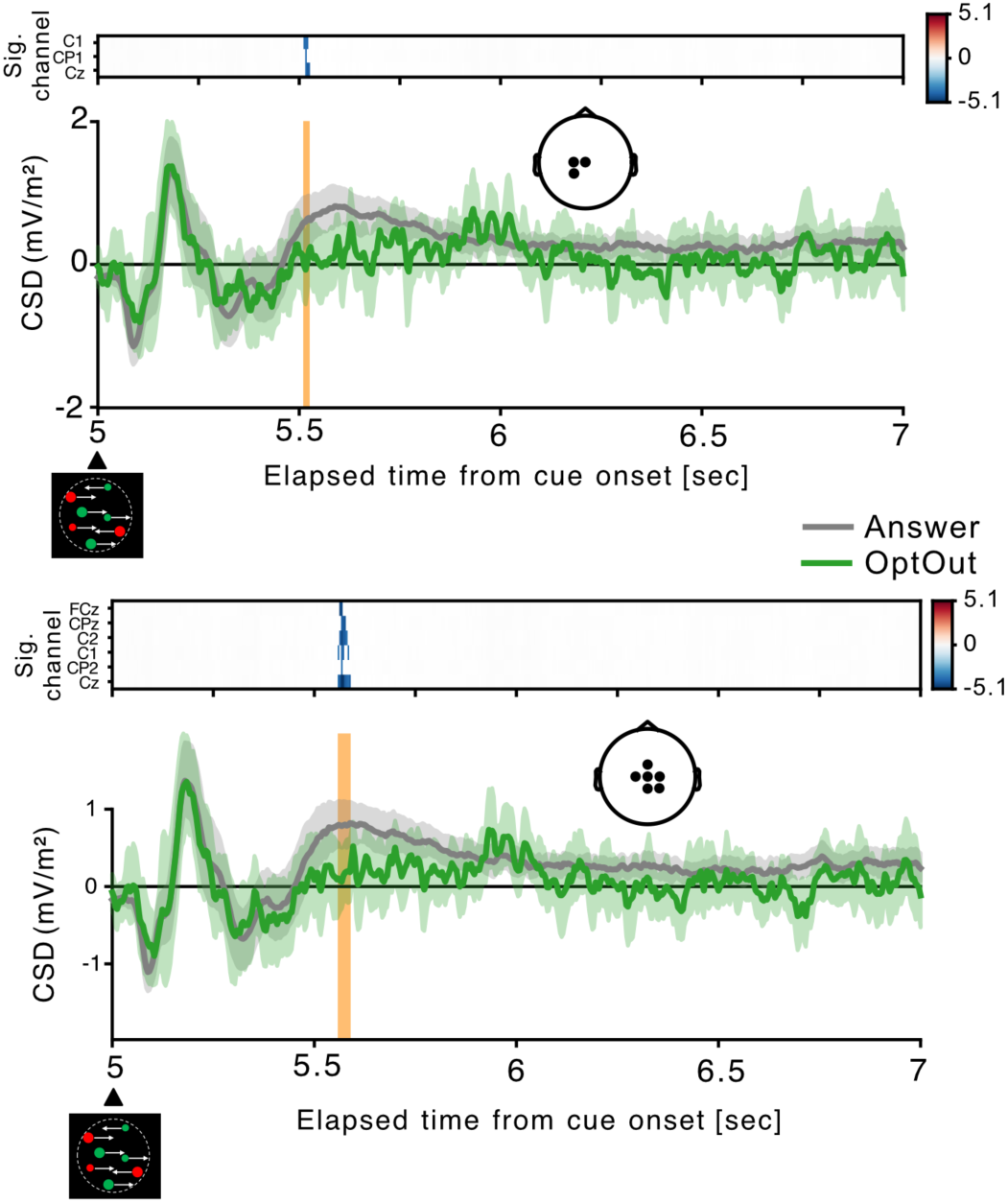
Changes in CPP components during decision-making. Based on cluster-based permutation testing, two significant EEG channel clusters (upper and lower panels) and corresponding time windows were identified. The channel clusters are shown in the topoplots, and the time windows are indicated by the orange vertical shaded lines. The CPP component was significantly greater in the Answer trials than in the Opt-out trials approximately 5.5 s after cue onset (i.e., 0.5 s after the onset of random dot motion). The lines and shaded areas indicate the mean ± 95% confidence interval across the participants for each condition.

To characterize the dynamics of neural activity during decision-making (Answer vs. Opt-out), we analyzed changes in brain functional connectivity in terms of connection density. Connection density was quantified to assess the degree of large-scale functional network recruitment underlying perceptual decision-making. Here, we identified two distinct neural signatures (Figure 3). The lower-frequency bands (delta and theta) showed significantly higher connection density from 2.5 seconds before the participants made the answer decision. In contrast, the higher-frequency bands (alpha and beta) exhibited significantly higher connection density after the participants made the answer decision. Regarding the spatial organization of the brain networks, although the present findings were obtained at the EEG sensor level, the increased connection density was primarily observed in bilateral occipital regions and the right frontal and parietal regions. In contrast, no significant increase in connection density was detected during the Opt-out trials.

**Figure 3:**
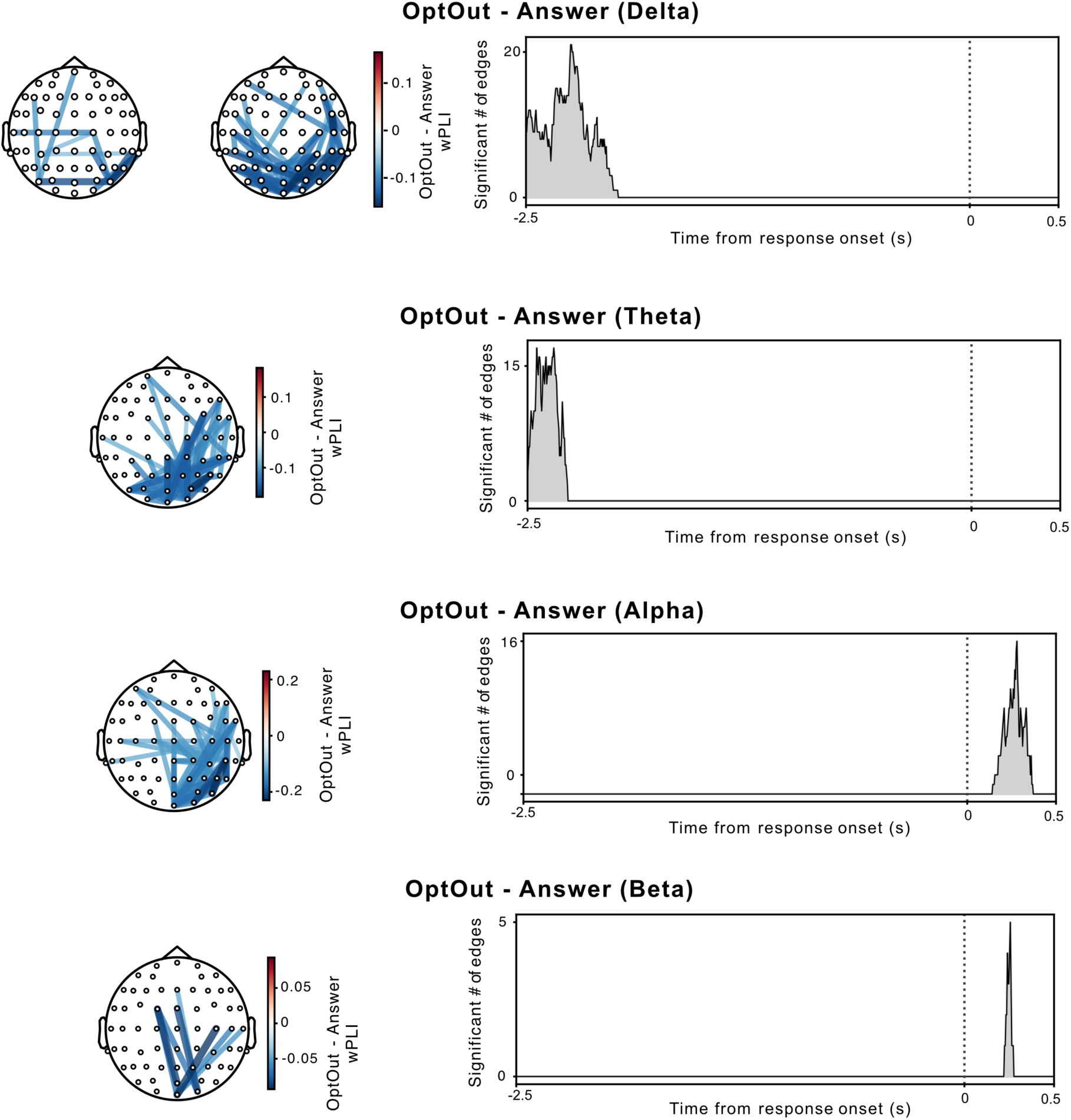
Brain network connectivity density around response onset (Answer vs. Opt-out). Significant differences in wPLI between the Opt-out and Answer trials are overlaid on the topoplots for each frequency band. Blue lines indicate connections with higher wPLI during the Answer trials. The time course of the number of significant EEG connections during Answer trials is shown in the right panel. The time axis was aligned to the keyboard response onset (0 ms on the x-axis) for the binary classification of Answer versus Opt-out trials. The gamma band is omitted from the figure because no significant wPLI changes were detected.

Moreover, we investigated whether distinct brain networks were associated with correct and incorrect responses, based on the assumption that participants were more confident when they provided the correct answer. Figure 4 shows the connectivity density and its spatial and temporal dynamics. Significant changes in connectivity density were observed across all frequency bands except the delta band. In the correct trials, significantly higher connectivity density emerged before participants made their answer decisions, and this connectivity gradually expanded from the occipital to the frontal regions. In addition, the parietal regions exhibited higher connectivity density in the theta band. In contrast, no significant increases in connectivity density were observed in the incorrect trials. Taken together, these findings demonstrate that the temporal evolution of large-scale functional connectivity differed according to behavioral outcome. Robust connectivity recruitment was observed during Answer and correct trials, whereas no significant network recruitment was detected during Opt-out or incorrect trials.

**Figure 4:**
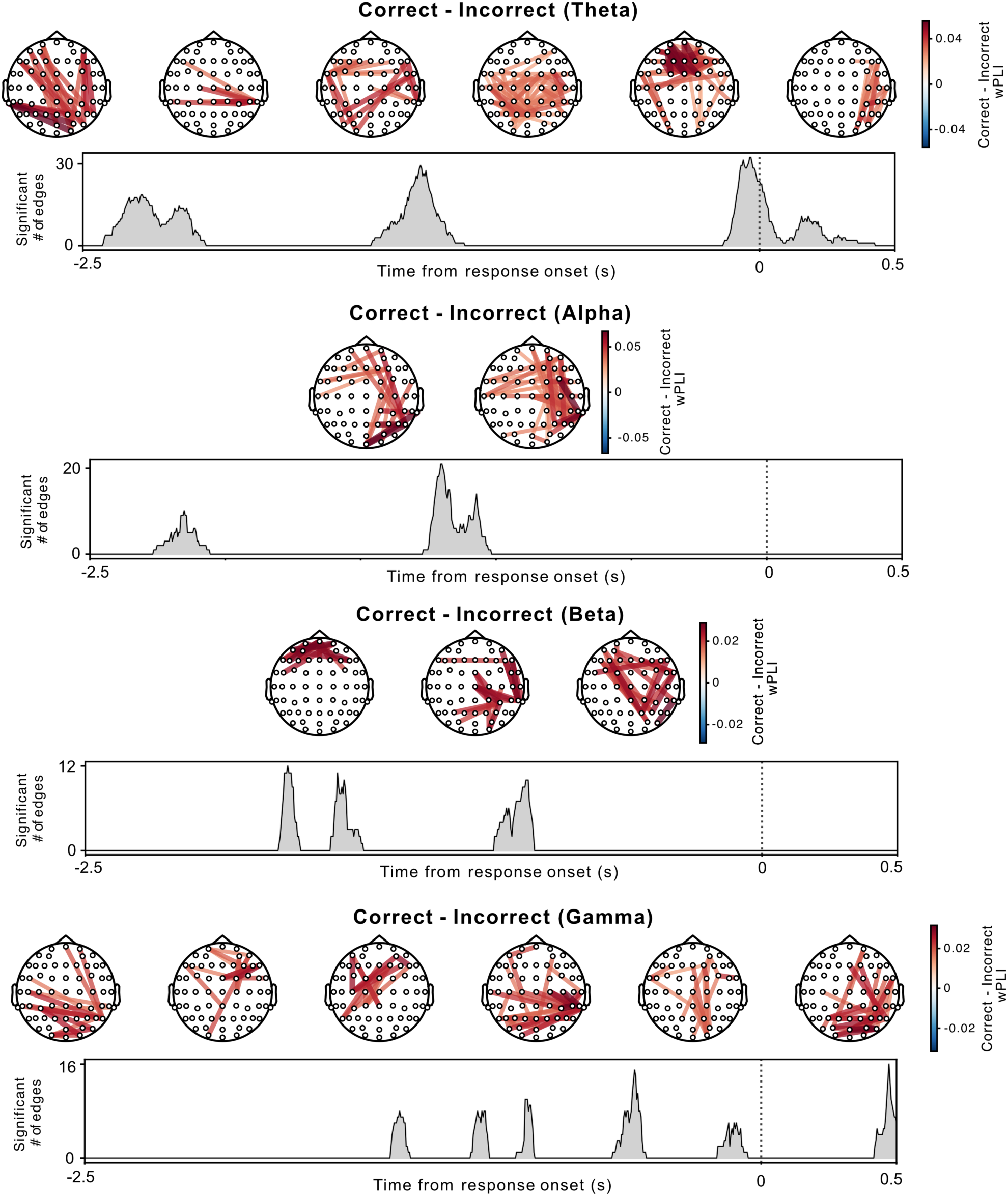
Brain network connectivity density for the correct and incorrect trials. Significant differences in wPLI between the correct and incorrect trials are overlaid on the topoplots for each frequency band. Red lines indicate connections with higher wPLI when participants responded correctly. The time course of the number of significant EEG connections for the correct trials is plotted alongside each topoplot. The time axis was aligned to the keyboard response onset (0 ms on the x-axis) for the binary classification of Answer versus Opt-out trials. The delta band is omitted from the figure because no significant wPLI changes were detected.

## Discussion

Using a novel approach in which perceptual uncertainty was parametrically manipulated and opt-out choices were included, we found that EEG-based functional connectivity patterns were associated with the emergence of confidence during perceptual decision-making. From a behavioral perspective, perceptual uncertainty was successfully manipulated by increasing the number of cues, resulting in systematic changes in reaction times and task success rates. As shown in Figure 1A, the task structure used in the present study required two separate responses (Choice response and Answer response) to assess the relationship between confidence and decision-making. Choosing to answer (i.e., challenging the given task) and responding correctly under conditions of uncertainty suggest that participants maintained high confidence while performing the task despite the uncertain environment. Our main behavioral measure was reaction time, which reflects multiple cognitive processes involved in decision-making. We found that perceptual uncertainty affected the time required to reach a decision, rather than the decision itself, and longer reaction times were associated with both opting out and incorrect responses. This indicates that perceptual uncertainty mainly affected the time course of decision formation, as reflected by prolonged reaction times, rather than exclusively altering the final decision outcome. As mentioned above, CPP serves as an electrophysiological marker of the accumulation of decision-relevant evidence toward a decision bound (Dou et al., 2024; O’Connell et al., 2012). Previous studies repeatedly reported larger CPP amplitudes co-occurring in trials with higher confidence ratings (Gherman & Philiastides, 2015, 2018; Herding et al., 2019; Rausch et al., 2020; Squires et al., 1973; Zakrzewski et al., 2019; Kelly & O’Connell, 2013). We further found an additional noteworthy result. The CPP amplitudes were significantly greater in the Answer trials than in the opt-out trials, whereas opt-out decisions reflect insufficient evidence accumulation (Figure 2). These findings were obtained through a whole-channel analysis without predefined channels of interest, providing data-driven evidence that the observed CPP modulation was independent of a priori channel selection. Additionally, based on our findings, the opt-out trials may not simply reflect low confidence, but rather a strategic termination of evidence accumulation before sufficient evidence reaches the decision bound. Thus, the distinction between the Answer and opt-out trials appears to emerge primarily from differences in the evidence accumulation process rather than from differences in the final decision outcome. We offer new possibilities that incorporating the opt-out choice can serve as an explicit behavioral index of uncertainty during decision formation, complementing conventional confidence ratings obtained after decisions.

Our behavioral and CPP findings empirically demonstrate that our experimental paradigm enables a more detailed investigation of the dynamic orchestration of brain activity across distributed cortical networks underlying confidence under perceptual uncertainty. A previous fMRI study showed that the BOLD (Blood-Oxygen-Level-Dependent) responses were increased with subjective confidence in the striatum, lateral orbitofrontal cortex, ventral anterior cingulate cortex, anterior middle frontal gyrus, amygdala, hippocampus, visual association areas, supplementary motor area, dorsomedial prefrontal cortex, inferior frontal gyrus, anterior insula, and frontal operculum (Gherman & Philiastides, 2018). These BOLD responses suggest that confidence depends on coordinated interactions among widely distributed brain regions, rather than being represented solely within local brain areas. To characterize the dynamic orchestration of neural activity at a higher temporal resolution than afforded by BOLD responses, we collected multichannel EEG data and estimated phase-based functional connectivity using the weighted phase lag index (wPLI). In the present study, we specifically focused on neural connection density (i.e., the number of connections over a specific temporal period), as increased connectivity may reflect enhanced network integration or compensatory recruitment of distributed neural resources (Finc et al., 2017; Schedlbauer et al., 2014; Tomasi & Volkow, 2010; Van Wijk et al., 2010). We found that network density significantly increased in the Answer trials within lower-frequency bands before the choice response. Interestingly, increased connectivity density emerged after the Answer response in higher-frequency bands (Figure 3). Regarding topological characteristics, increased network density was observed in interhemispheric occipital connections and connections between the right occipital and frontal areas during Answer trials. A plausible interpretation for these findings is that interhemispheric occipital connectivity may reflect enhanced integration of bilateral visual information required to accumulate sensory evidence during perceptual decision-making and connectivity between occipital and frontal regions may reflect enhanced communication between sensory processing regions and higher-order decision-related regions during confidence formation. Answer trials may reflect enhanced communication between sensory and decision-related regions. Such increased connectivity may facilitate the integration of visual evidence and support the formation of confident decisions. This interpretation is also consistent with the increased CPP amplitude observed in the Answer trials. In addition, our findings of the connectivity density exhibited a predominance of right-hemispheric connectivity. This is in line with previous studies showing that the right anterior prefrontal cortex plays a role in metacognition monitoring (Fleming et al., 2010) and that confidence computations occurred at the right dorsolateral prefrontal cortex (Xue et al., 2023). The right frontopolar cortex is important for the reliable rating of confidence in past short-term recognition memory performance (Yokoyama et al., 2010). Taken together, we therefore suggest that the predominance of right-hemispheric connectivity during Answer trials may reflect enhanced recruitment of right-lateralized frontoparietal networks involved in monitoring decision reliability and forming confidence. When participants committed to a perceptual decision, increased communication between the occipital sensory-related regions and frontal decision-related regions may have supported the integration of sensory evidence with internal confidence estimates.

By comparing correct and incorrect responses, we further investigated the neural mechanisms underlying confidence formation under perceptual uncertainty. As described above, increased connectivity density emerged in correct trials across all frequency bands but was not observed in incorrect trials (Figure 4). Interestingly, although the temporal profile of connectivity density was aligned with the choice time (i.e., Answer or Opt-out), distributed brain regions in correct trials had already established functional connections before the answer decision was made. Given that such connectivity enhancement was absent in incorrect trials, correct decisions may be characterized by the early recruitment of distributed neural networks that support the integration of sensory evidence and the formation of reliable confidence. Consistent with this interpretation, previous animal studies have shown that decision-related neural activity in the lateral intraparietal cortex reflects accumulated evidence before behavioral responses are initiated (Kiani & Shadlen, 2009). Furthermore, a human fMRI study has demonstrated that distributed prefrontal and parietal activity contain information predictive of upcoming decisions before conscious awareness emerges (Soon et al., 2008). These findings, together with previous evidence, suggest that increased whole-brain connectivity preceding the decision-making may reflect the establishment of a distributed neural state supporting evidence integration and confidence formation.

The present study has an important caveat that should be considered when interpreting our findings. Although we adjusted task difficulty and identified parameters using another independent cohort, we were unable to fully minimize individual differences in behavioral performance, such as the number of opt-out trials. This limitation in task design may have limited the generalizability of the EEG findings. In future studies, individual task difficulty should be determined before data collection.

In conclusion, we developed a novel perceptual decision-making paradigm that parametrically manipulated perceptual uncertainty and incorporated opt-out choices as an explicit behavioral index of uncertainty. Our findings demonstrate that confidence formation under perceptual uncertainty was associated with dynamic changes in functional connectivity across distributed brain networks, particularly between occipital and frontal regions. These results provide novel evidence that confidence emerges through the dynamic integration of sensory evidence across distributed neural networks. Moreover, we were unable to detect specific neural features associated with decision uncertainty (i.e., opt-out or incorrect trials), suggesting that neural activity may not have stabilized into a distinct functional network state, reflecting continued transitions among competing neural states during decision formation.

## Data availability statement

The data and code that produce the findings in this study are available from the corresponding author upon reasonable request.

## Acknowledgements

This work was supported by JSPS KAKENHI Grant Number JP24K02842 to K.U. and JST SPRING, JPMJSP2171 to T.O.

